# Shared viral burdens: Evidence of active Usutu virus circulation and multi-arbovirus exposure in migrant and resident birds at wintering locations in Nigeria

**DOI:** 10.1101/2025.09.08.674803

**Authors:** Nnomzie C Atama, Joy A Ishong, Cora M Holicki, Yahkat Barshep, Felicity D Chandler, Anne van der Linden, Marion P G Koopmans, Reina S Sikkema

## Abstract

**Background:** West Nile (WNV), Usutu (USUV), and Sindbis (SINV) virus were initially detected in the African region, and subsequently across temperate regions where they were absent. Wild birds are primary reservoirs for these arboviruses and are considered major contributors to their global spread through seasonal migration. To understand the transmission dynamics of arboviruses in wild birds and the potential of migratory birds to spread the viruses at an intercontinental scale, we investigated arboviral infections and exposures in African resident and Palearctic migratory birds at wintering locations in Nigeria.

**Methodology/Principal Findings:** Oropharyngeal- and cloacal swabs, feathers and blood were collected from resident and migratory birds at two wintering locations (Amurum and Ngel-Nyaki Forest Reserves). Swabs and feathers were tested using RT-PCR for WNV, USUV and SINV, and blood with ELISA and FRNT_90_ or PRNT_80_ for antibodies. 573 birds were sampled between 2021 to 2024 across months coinciding with arrival and departure of migratory birds. USUV RNA was detected in 2.6% of feathers including a positive Icterine warbler and a garden warbler sampled prior to spring migration. None of the swabs was positive for viral RNA but neutralizing antibodies to WNV and USUV were detected in 4.5% of birds. SINV antibodies were also found in 34.1% of birds sampled across the wintering locations.

**Conclusions/Significance:** Our findings showed that migratory birds can become infected with USUV, and potentially with WNV and SINV during their overwintering periods in Africa and highlighted a wider arbovirus risk in Nigeria. In addition, detections of viral RNA in feathers, but not swabs, suggest feathers may be a suitable matrix for surveillance in the absence of a reliable cold chain. The overall detections in wild birds at these locations highlight the need for further surveillance to define the epidemiology and public health risks of these arboviruses in the region.

**Synopsis:** Vector-borne viruses like West Nile, Usutu, and Sindbis, were once limited to parts of Africa, but now appear in Europe and other temperate regions. Migratory birds are believed to help spread these viruses across continents. To understand the role of birds in this, we tested resident and migratory birds in Nigeria for evidence of infection. Between 2021 and 2024, we sampled over 570 birds during their winter stay. We found genetic material of Usutu virus in feathers of six birds including two migratory birds just before their return to Europe. While swab tests did not detect active infections, antibodies to these viruses, which indicate past exposure, were found in several birds, especially for Sindbis virus. Our findings show that migratory birds can pick up and possibly carry these viruses across continents. We also found that feathers may be a practical tool for virus detection in areas without reliable refrigeration, supporting easier and broader surveillance of these viruses in Africa and possibly beyond.

## Introduction

Several mosquito-borne arboviruses, including viruses in the *Flaviviridae* and *Togaviridae* families, are primarily maintained in enzootic transmission cycles between wild birds and mosquitoes [1]; however, others predominantly circulate among humans or mammals [2]. Wild birds have long been identified as reservoir and amplifying hosts for several of these arboviruses, including flaviviruses like West Nile virus (WNV) and Usutu virus (USUV), and alphaviruses like Sindbis virus (SINV) and Western Equine Encephalitis virus (WEEV). Several of these viruses are of veterinary and/or public health significance. For example, WNV has caused many outbreaks in humans and animals [3]. Also, USUV can cause morbidity and mortality in animals, but is considered less pathogenic in humans [4].

Following initial isolation of USUV from field caught *Culex neavei* mosquito in Southern Africa in 1959 [5], the virus was subsequently detected in mosquitoes and vertebrate hosts, mainly birds, across several African countries including Senegal, Central African Republic, Nigeria, Uganda, Burkina Faso, Cote d’Ivoire, and Morocco [6–8]. In Europe, USUV was first detected in 2001 in Vienna, Austria, where it was identified as the cause of mortality in wild birds [9].

Phylogenetic analysis of USUV genomic sequences across Europe revealed similarity and transmission links between USUV strains in Africa and Europe indicating multiple independent introductions through migratory avian hosts [10].

WNV was first isolated in 1939 from the blood of a febrile patient at the West Nile district of Uganda [11]. Since then, the virus has caused sporadic outbreaks in many parts of the world. Genomic evidence shows that some outbreaks in Europe were seeded through migratory birds coming from Africa [12]. WNV has caused high mortality in birds in America, however its impact in birds across Europe has been minimal [13].

SINV is an alphavirus that is transmitted mainly by *Culex* mosquitoes such as *Culex neavei*,*Culex pipiens sensu stricto (s.s.),* and *Culex torrentium* [14,15]. It was first detected in mosquitoes in Egypt in 1952 [16], and has since been found in birds and humans in South Africa [17] and across Northern Europe, and recently in other European countries including Spain, Germany, Romania, and the Netherlands [18–21]. SINV has birds as primary amplifying hosts, and infections in humans often result in mild disease characterized by fever and rash, and sometimes arthralgia, with symptoms persisting for months in a considerable percentage of patients [17].

Within Africa, arboviruses like USUV, WNV, and SINV are maintained in circulation between birds and *Culex* mosquitoes, and infections in local host species seem to cause only minimal fatality. So far, there is scanty evidence for USUV and WNV circulation in birds and humans in Nigeria [6,22–24]. In 1972, USUV was isolated from passerine bird species including Little greenbul (*Eurillas virens*), Piping hornbill (*Bycanistes fistulator*), and Kurrichane thrush (*Turdus libonyana*) [6] (Fig 1C). Thereafter, serological evidence for WNV was documented in domestic birds and humans across Nigeria [24–26]. SINV, on the other hand, was only recently detected for the first time in Nigeria in African thrushes (*Turdus pelios*) [27].

**Fig 1.**
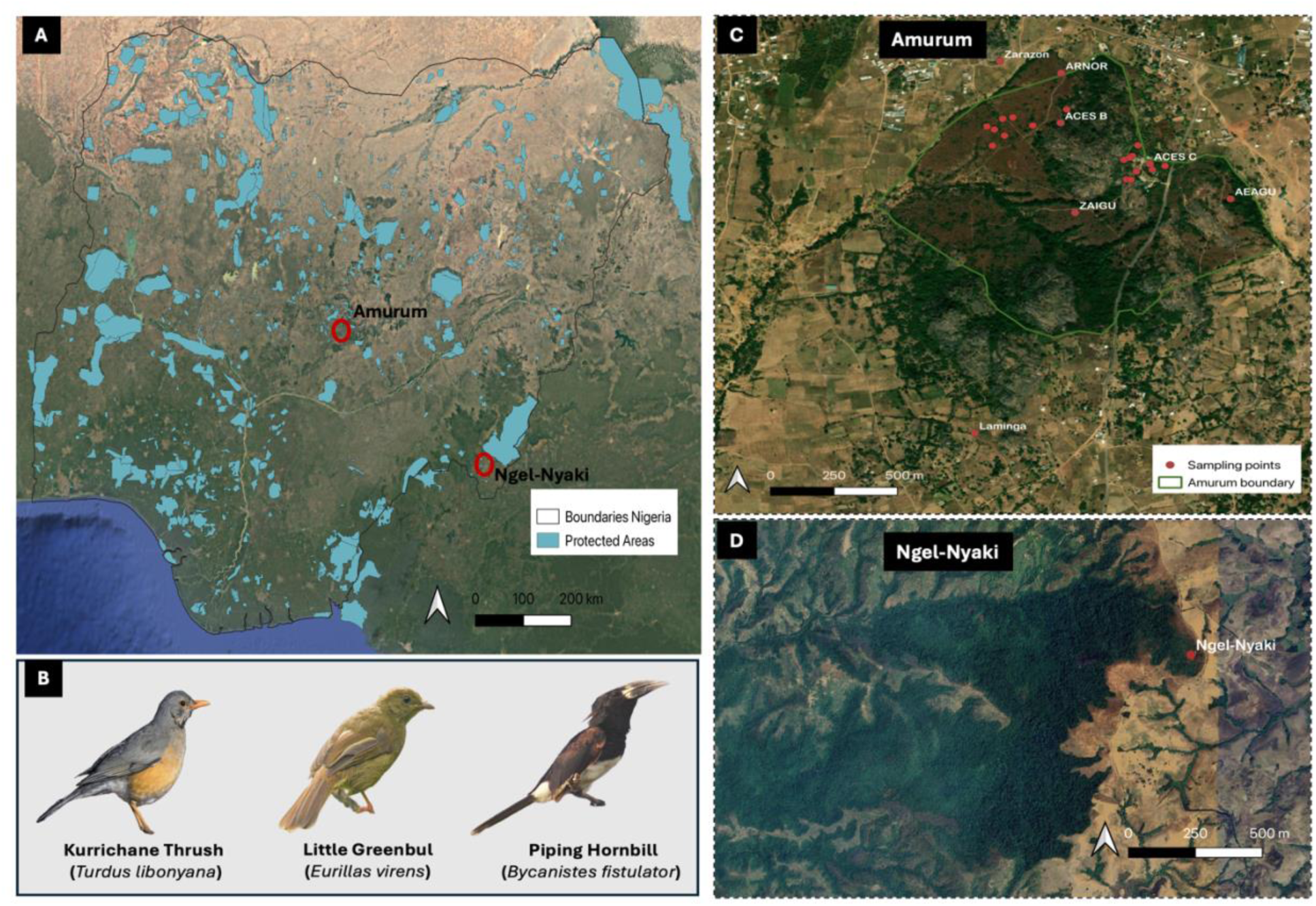
Sampling sites across two overwintering locations in Nigeria. 1**A**. Nigeria with locations of protected areas (light blue patches) [35] and sampled locations (red circles) – Amurum Forest Reserve (Amurum) and Ngel-Nyaki Forest Reserve (Ngel-Nyaki). 1**B.** *In 1972, Usutu virus was isolated from three species – Kurrichane thrush, little greenbul, and piping hornbill, at an unspecified location in Nigeria* [6,36]. Bird photos adopted from ebird (https://ebird.org/species/). 1**C**. **Amurum,** sampling was conducted at constant effort sampling (CES) sites within the reserve including CESB, CESC, Zainab’s Gully (ZAIGU), Eastern Gully (AEAGU), North of the Reserve (ARNOR), and at outside locations bordering the reserve - Laminga village (Laminga) and Zarazon. **NB**. mist-netting points indicated in red. 1**D**. Ngel-Nyaki, sampling was conducted at one location (section of Ngel-Nyaki on map).

As a tropical location with diverse habitats and known Important Bird Areas (IBAs), Nigeria is home to a great diversity of birds including endemic species. It serves as an overwintering location for about 253 migratory birds [28]. Several Palearctic migratory bird species that are known reservoirs for arboviruses undertake seasonal migration to overwinter in Nigeria and other parts of Africa [29,30], and their role in the intercontinental spread and emergence of arboviruses is yet to be fully resolved. Therefore, understanding the infection and exposure status of these groups of birds at their overwintering and stopover locations will shed more light on their role in transmission across bird flyways and on the evolution of the virus. Here, we aimed to identify and characterize the circulation and transmission of arboviruses including WNV, USUV, and SINV in African resident and Palearctic migratory birds at two overwintering IBAs in Nigeria – Amurum Forest Reserve (Amurum) and Ngel-Nyaki Forest Reserve (Ngel-Nyaki).

## Methods

### Ethics statement

The handling and sampling of birds was carried out under protocols and permits approved by the Scientific Committee of the A P Leventis Ornithological Research Institute (APLORI), University of Jos, Nigeria. A material transfer agreement (MTA) approved by the Nigerian Federal Ministry of Environment with the FMENV MTA Reference: FOR/ABS PERMIT/12/1 permitted transfer of samples between APLORI and Erasmus Medical Center.

### Study sites

The study was conducted in Nigeria at two protected areas – within and around the vicinity of Amurum Forest Reserve (Amurum) and at Ngel-Nyaki Forest Reserve (Ngel-Nyaki). Amurum is in the North-Eastern part of Plateau State in North-Central Nigeria (9°53′N, 8°59′E) covering an area of 300 hectares and composed of granitic outcrops and patchy grasslands with interspersed gallery forests [31]. Ngel-Nyaki, however, is a montane forest area in the Mambila plateau located 60 km west of Gashaka-Gumti National Park (7 .08° N, 11.13 °E). It extends to about 45 km^2^ with 7 – 7.2 km^2^ of dense montane forest and an 85% shrubland cover. Its wet season lasts from March to November with an average annual rainfall of about 2,000 mm [32] (Fig 1C & 1D). Both locations are protected areas rich in biodiversity and avifauna and are recognized as IBAs according to Birdlife International [28]. They both also serve as breeding locations for resident and intra-African migratory birds, and as overwintering locations for Palearctic migratory birds [28] that mostly arrive the wintering locations around September/October and depart in spring (March/April) [33,34].

### Bird sampling and sample collection

As part of APLORI’s Constant Effort ringing scheme [37] conducted across Constant Effort Sites (CES) in and around Amurum and at Ngel-Nyaki, wild birds including African resident and Palearctic migratory birds, were mist-netted, ringed, and sampled (Fig 1C & 1D). In Amurum, sampling was conducted across dry and wet seasons, between March - December 2021, February - April 2022, and April - December 2024. At Ngel-Nyaki, however, samples were only collected once, in March of 2022. Birds were sampled across seven sites within and around Amurum, and at one location in Ngel-Nyaki (Fig 1D). Sampling was conducted to tally with periods spent by migratory birds at the wintering locations, based on over 20 years of observational and ringing data on migratory birds through APLORI Bird Ringing (APBRING) Scheme [37]. Birds caught were identified to species, aged (from plumage as years relative to hatch year), and sexed when possible. Ringed birds were sampled by collecting swabs (oropharyngeal and/or cloaca) in 1.5 mL Invitrogen RNA Later ^TM^ (ThermoFisher Scientific, Vilnius, Lithuania). For smaller birds, only oropharyngeal swabs were collected, whereas in the larger birds, cloacal swabs were also taken. Wing or breast feathers were collected, and blood collected by brachial venipuncture when possible. Biometric measures including wing length, body mass, fat score, and moult score were taken. In 2024, only feather samples were collected from sampled birds at Amurum. All samples were first shortly stored under +4°C before being transferred to -80°C and later shipped on dry ice to the WHO arbovirus reference laboratory at Viroscience Department of Erasmus Medical Center in Rotterdam, The Netherlands for molecular and serological analyses.

### Molecular analysis

Swab samples were prepared by first transferring swabs from RNA later into 2 mL sample tubes containing virus transport medium, vortexed, and centrifuged at 10,000 xg for 1 minute. Total nucleic acid (tNA) was extracted by eluting 400 μL of swab material in 600 μL of Roche external lysis buffer and processed using an MP96 MagNA pure LC instrument (Roche Diagnostics GmbH, Mannheim, Germany). The larger wing feather samples were processed by splitting 1cm of the lower half of the feather calamus and homogenizing in 350μL Roche tissue lysis buffer plus ¼” ceramic beads using a FastPrep-24 5G tissue Magnilizer (M.P. Biomedicals, LLC, California, USA). The smaller breast feathers were, however, homogenized whole at 60 sec; 6.5m/s as previously described [38].

Extracted RNA materials (tNA) were tested on screening and confirmatory RT-PCRs for WNV, USUV, TBEV, JEV, and SINV targeting different genomic regions of each of the viruses with primers and probes as described previously [39]. RNA from swabs were tested in pools of five, while feather samples were tested individually. Samples were considered positive only if they had cycle threshold (C_T_) values < 40 in both the screening and confirmatory PCR assays. When C_T_-values of positive samples were too high to sequence (> 32), blood clots of positive birds, which were obtained after separation of serum, were also tested. Adapting from a previously described protocol [40], a pinhead-size of blood clot was added to 600 μL tissue lysis Buffer and 6 μL β-mercaptoethanol with 5 mm ceramic beads and homogenized at 4 m/s for 20sec using the FastPrep-24 tissue Magnalizer. Then 50 μL of the supernatant was eluted in 150 μL binding/lysis buffer and 400 μL PBS to extract RNA based on a manual column extraction protocol using High Pure RNA Isolation kit (Roche Diagnostic GmbH, Mannheim, Germany) according to manufacturer’s instruction. In general, samples that were positive on both screening and confirmatory PCRs were also tested on hemi-nested and nested pan-PCRs designed for flaviviruses and alphaviruses targeting conserved regions of NS5 gene and NSP4 gene, respectively, as previously described [41,42] . The products of the pan-PCRs were then submitted for sequencing using a Sanger sequencing approach.

### Serological analysis

Blood samples were centrifuged at 10,000 xg for 5 minutes to collect serum. Most of the blood samples were stored longer before centrifugation which resulted in impure sera with red blood cell content. Collected supernatant were stored under -80°C until they were analyzed.

#### ELISA

All serum samples were screened for binding antibodies to the domain III of the E protein of WNV using a commercial blocking ELISA (*INGEZIM West Nile COMPAC – Eurofins Technologies, Madrid, Spain*) according to the manufacturer’s instructions, where an inhibition percentage (IP) ≥ 40% was termed positive, IP > 30% or < 40% as doubtful and ≤ 30% as negative. The ELISA kit has the potential to detect cross-reactivity with USUV and was thus used to also capture antibodies to USUV if present [43,44]. Samples reactive on ELISA (positive or doubtful) were further tested for neutralizing antibodies using a focus reduction neutralization test (FRNT) to differentiate positive from doubtful ELISA results and classify reactivity as either to WNV or USUV.

#### WNV-USUV FRNT

The ELISA positive samples were tested for WNV and USUV neutralizing antibodies using in-house validated assays (WNV- and USUV-FRNT) that assess presence of neutralizing antibodies based on a 90% reduction in foci (cluster of infected cells) on a confluent monolayer of Vero cells for WNV and USUV, respectively [45]. Initial testing demonstrated that the FRNT read-out was hampered by cell debris from the poor-quality sera. Therefore, an additional washing step with subsequent addition of infection medium (DMEM supplemented with FBS) was added to the protocol after a 2-hour incubation step of the serum-virus mix on the cell monolayer. The plates were then incubated for 24 hours at 37°C following standard protocol. A sample was defined as positive for WNV or USUV if it showed neutralization at titer ≥ 80 (for WNV) or ≥ 160 (for USUV) and a 4-fold difference in titer between WNV and USUV or vice-versa.

#### SINV PRNT

Due to limited serum volumes, only samples negative in the WNV ELISA, which still had serum left, were tested for neutralizing antibodies to SINV (strain 09-M-991-1 from Sweden, Accession number: KF297653) using a plaque reduction neutralization test (PRNT) on a Vero ATCC CCL-81 cell line [18,46] Samples that were positive in the pre-screening (1:20 dilution) were serially diluted from 1:20 to 1:640. Samples were dimmed positive if they showed 80% reduction of plaque counts compared to a virus control in the pre-screening and end-titration. Due in part to the poor quality of the sera, individual samples were pre-screened or end-titrated a second time, and the average titer was calculated. Samples that were too haemolytic for the read-out were excluded.

### Data Analysis

Data management, analysis, and visualization of graphs were done using Microsoft Excel v16.95.4 and RStudio v2024.12.1.563 [47]. SINV Seroprevalence was estimated as proportion of positive individuals among the total tested, with exact binomial 95% confidence intervals (CIs) calculated using the Wilson score method [48]. We fitted a mixed-effects logistic regression model using the lme4 package in R, with SINV seropositive status (positive/negative) as dependent variable. Migratory status and season were included as fixed effects, while species and sampling location were treated as random intercepts to account for clustering and reduce collinearity. Performance package in R was used to assess model performance [49].

## Results

573 birds of 59 species were sampled in 2021, 2022, and 2024. In 2021 and 2022, wild birds were sampled for swabs (oropharyngeal and/or cloaca), feather, and blood, whereas in 2024 only feather samples were collected. Most samples were collected from birds at Amurum and only 12 of the 573 birds were sampled at Ngel-Nyaki (Fig 1D). 51.1% (293) of the sampled birds were migratory birds comprising of 17 species, while the others were African resident birds.

52.5% (301) of the samples were collected during the wet seasons of the sampling years (between late April to early October) when mosquito abundance is highest [50]. Swabs were collected from 464 birds, feathers from 229, and blood from 264 birds.

### Molecular detections

All swabs were negative for arboviruses in RT-PCR. However, USUV RNA was found in feathers of six birds at a prevalence of 2.6% (6/229; 95% CI: 1.2 - 5.6). None of the feathers were positive for WNV, TBEV, or JEV. A feather sample from a Spotted flycatcher (*Muscicapa striata*) tested positive for SINV on a first PCR (C_T_ 35) but could not be confirmed using a second PCR that targeted a different region of the SINV genome. None of the blood clots from the USUV positive feathers tested positive for USUV RNA. The six USUV positive birds included three African resident birds (two African thrushes; *Turdus pelios* and a Black-necked weaver; *Ploceus nigricollis*), and three Afro-Palearctic migrant birds (Tree pipit; *Anthus trivialis*, Icterine warbler; *Hippolais icterina*, and Garden warbler; *Sylvia borin*). The USUV positive African thrushes, Black-necked weaver, and Tree pipit were sampled during the dry season (between December 2021 and February 2022). The positive Icterine warbler and Garden warbler were however sampled during the wet season; in April 2022 and October 2024, respectively. The C_T_-values of the positive birds ranged between 34.9 – 37.1 (median C_T_ 35.6). Except the positive garden warbler, which was sampled at Laminga, a village bordering Amurum, all other RNA positive birds were trapped and sampled within the reserve, at the Constant Effort Site B (ACESB) – a location where 57.9 % (332) of the total birds were sampled (Fig 1C).

All sequencing attempts failed due to low viral loads in positive samples and no genomic sequences were generated from any of the USUV positives.

### Serologic detections

46 out of 264 (17.4%) serum samples were reactive on the ELISA, with 35 positives (IP ≥40%) and 11 doubtful (IP >30% and <40%). 25 of the 46 reactive samples (18 positives & 7 doubtful) had insufficient sera volume remaining and were not tested for neutralizing antibodies on FRNT.

An overall 4.5% (12/264; 95% CI: 2.6 - 7.8) of the birds had neutralizing antibodies to flaviviruses. Four out of 21 ELISA positive/doubtful samples were confirmed to have neutralizing antibodies to either WNV (n = 1) or USUV (n = 3). Eight other samples (8/21) which could not be differentiated based on 4-fold titer difference (n = 3) or that were neutralizing on either the WNV or USUV FRNT but had insufficient sera volume to test on both assays (n = 5) were classified as Orthoflavivirus positive (FLAVI) (Table 1; Fig 2). The WNV- and USUV-antibody positive birds were all African thrushes (resident bird species) sampled in the wet season of 2021. The FLAVI positives, however, included resident birds (three African thrushes, a blackcap babbler; *Turdoides reinwardtii*, a yellow-crowned gonolek; *Laniarius barbarus*, and a laughing dove; *Spilopelia senegalensis*) and migratory birds (two garden warblers). All the positive birds were sampled across four CES (ZAIGU, AEAGU, ACESB and ACESC) within Amurum, except one FLAVI-positive garden warbler that was sampled at Ngel-Nyaki (Table 1; Fig 2).

**Fig 2:**
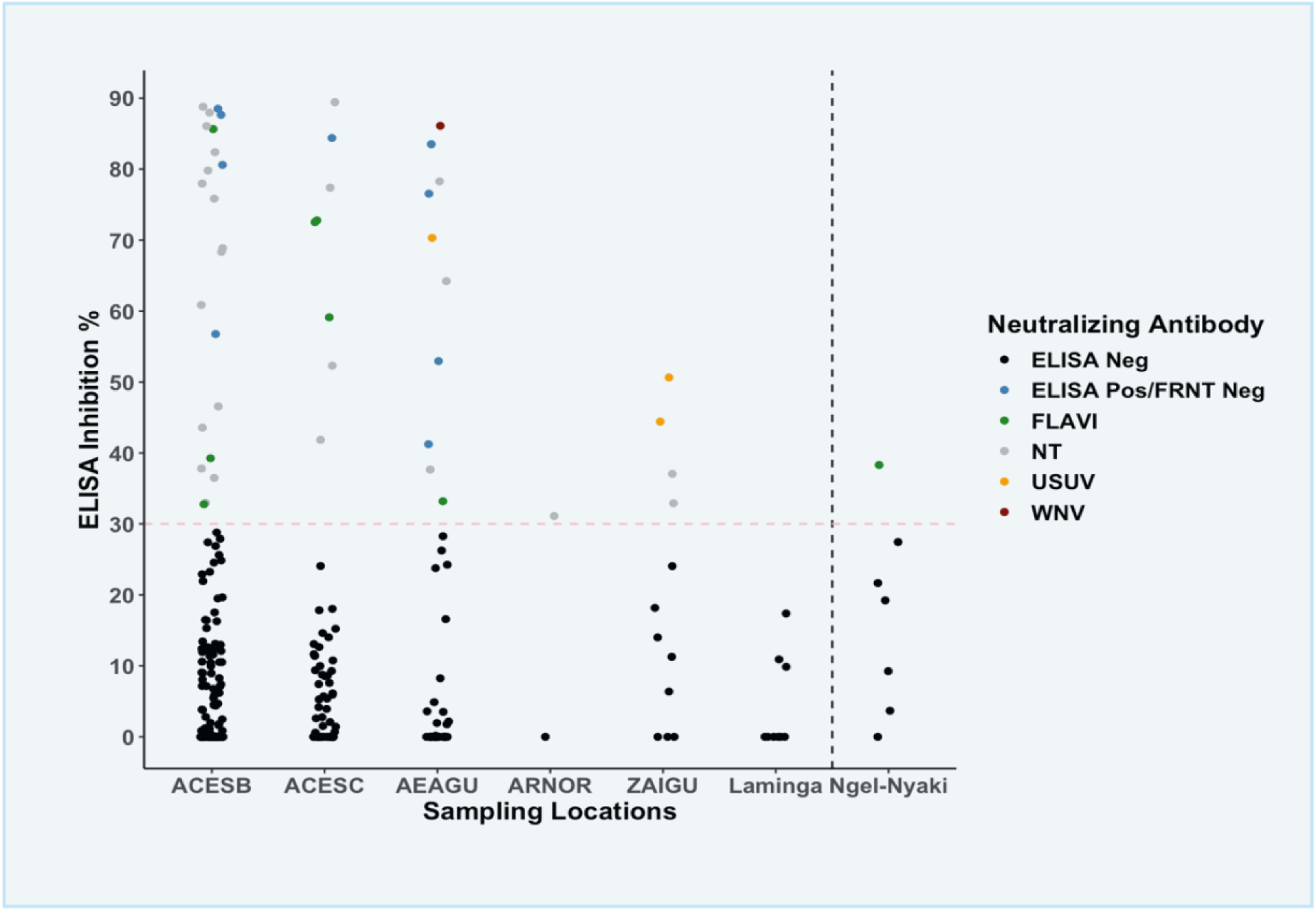
Locations of WNV, USUV, and Orthoflavivirus (FLAVI) positive birds. Virus-specific antibody responses were confirmed via FRNT_90_ for WNV (red), USUV (orange), and FLAVI (green) from ELISA reactive samples (ELISA Inhibition % ≥ 30). ELISA reactive samples with insufficient volume could not be further tested on FRNT_90_ (NT).

**Table 1:** ELISA positive samples across Afro-Palearctic migrant and African resident birds confirmed on FRNT for neutralizing antibodies to WNV, USUV, or Orthoflavivirus (FLAVI). NB.^☨^ are samples that were seropositive on the USUV FRNT but could not be retested with washing due to insufficient volumes and were thus marked as Orthoflavivirus-positive (FLAVI).

| Bird ID | Species | Ring No. | Migratory Status | Sampling Site | Sampling Year | Blocking ELISA |  | FRNT<sub>90</sub> |  |  |
| --- | --- | --- | --- | --- | --- | --- | --- | --- | --- | --- |
|  |  |  |  |  |  | Inhibition Percentage (IP%) | Results | WNV Antibody Titers | USUV Antibody Titers | Confirmed Antibody Status |
| 22NG0286 | African Thrush | 4A61014 | Resident | ZAIGU | 2021 | 50.6 | POS | 160 | 640 | USUV |
| 22NG0287 | African Thrush | 4A61013 | Resident | ZAIGU | 2021 | 44.4 | POS | 10 | 640 | USUV |
| 22NG0260 | African Thrush | 4A61020 | Resident | AEAGU | 2021 | 33.2 | DOUBTFUL | 160 | NT | FLAVI |
| 22NG0268 | African Thrush | 4A62984 | Resident | AEAGU | 2021 | 86.1 | POS | 320 | 20 | WNV |
| 22NG0297 | African Thrush | 4A61027 | Resident | AEAGU | 2021 | 70.3 | POS | 10 | 320 | USUV |
| 22NG0338 | African Thrush | 4A62267 | Resident | ACESC | 2021 | 59.1 | POS | 10 | 320 <sup>†</sup> | FLAVI |
| 22NG0370 | African Thrush | 4A61831 | Resident | ACESC | 2021 | 72.8 | POS | 10 | 320 <sup>†</sup> | FLAVI |
| 21NG014 | Blackcap Babbler | DH05223 | Resident | ACESB | 2021 | 85.6 | POS | 160 | NT | FLAVI |
| 21NG0076 | Garden Warbler | Y24390 | Migrant | Ngel- Nyaki | 2022 | 38.3 | DOUBTFUL | NT | 160 | FLAVI |
| 21NG0064 | Yellow-crowned Gonolek | 4A62656 | Resident | ACESC | 2022 | 72.5 | POS | 80 | NT | FLAVI |
| 21NG0015B | Laughing Dove | D96119 | Resident | ACESB | 2022 | 32.8 | DOUBTFUL | 20 | 320 <sup>†</sup> | FLAVI |
| 21NG0165 | Garden Warbler | AR8057 | Migrant | ACESB | 2022 | 39.3 | DOUBTFUL | 160 | NT | FLAVI |

Functional (neutralizing) antibodies were also detected in 45 of 132 (34.1%, 95% CI: 26.6 - 42.5) birds tested via SINV PRNT_80_. Antibodies to SINV were found in 23 bird species, representing eight migratory species groups and 15 resident species (Table 2). Besides two speckled mousebirds; *Colius striatus* (resident birds) and a blackcap; *Sylvia atricapilla* (a migratory bird) which were sampled at Ngel-Nyaki, all other SINV antibody-positive birds were sampled across different sites in Amurum. SINV antibody titers in both migratory and resident birds ranged between 20 to 320 (median titer 40). An estimated seroprevalence of 46.9% (95% CI: 30.6 - 63.9) was recorded in migratory birds, and 30.0% (95% CI: 21.8 - 39.7) in resident birds. In the mixed-effects logistic regression model, neither migratory status of the birds nor season showed statistically significant associations with SINV seropositivity. Resident birds had lower odds of SINV antibody detection compared to migratory birds (OR = 0.52, 95% CI: 0.19 – 1.47, p = 0.22), although this effect did not reach significance. Similarly, individuals sampled during the wet season, exhibited higher odds of seropositivity compared to those sampled during the dry season (OR = 1.61, 95% CI: 0.63 – 4.09, p = 0.32), but again, this difference was not significant. The wide confidence intervals indicate considerable uncertainty, likely due to limited sample sizes and variability among species and locations.

**Table 2:** Overview of SINV antibody-positive Afro-palearctic migrant and African resident bird species.

| **Species** |  |  |  |  | **SINV PRNT<sub>80</sub>** |
| --- | --- | --- | --- | --- | --- |
| **Paleartic Migrant** | **Sampling Sites** | **Seasons** | **Months** | **Years** | **Pos/Tested (~ %)** |
| Blackcap | Ngel-Nyaki | dry | Mar | 2022 | 1/2 (50) |
| Common Whitethroat | AEAGU | dry | Mar | 2022 | 1/2 (50) |
| European Nightjar | ACESB | wet | Apr | 2022 | 1/1 (100) |
| Garden Warbler | ACESB, ACESC | wet | Apr, Mar | 2021- 2022 | 4/6 (67) |
| Icterine Warbler | ACESB, ACESC | wet | Apr | 2021 | 2/2 (100) |
| Pied Flycatcher | ACESC | wet | Oct | 2021 | 2/5 (40) |
| Spotted Flycatcher | ACESB, ACESC | wet | Apr | 2021 | 3/8 (38) |
| **African Resident** |  |  |  |  |  |
| Adamawa turtle dove | ACESB | dry | Feb | 2022 | 1/12 (8) |
| African Thrush | ACESB, ACESC, AEAGU | dry, wet | Feb, Mar, Apr, Dec | 2021-2022 | 9/18 (50) |
| Common Bulbul | ACESC, AEAGU, ZAIGU | dry, wet | Apr, Mar, Dec | 2021, 2022 | 3/6 (50) |
| Mocking Cliff Chat | ACESC | wet | Oct | 2021 | 1/2 (50) |
| Moustached Grass Warbler | ACESC | wet | Apr | 2022 | 1/1 (100) |
| Northern Red Bishop | ACESB | wet | Apr | 2022 | 1/1 (100) |
| Orange-breasted Bushshrike | ACESC | dry | Feb | 2022 | 1/1 (100) |
| Senegal Coucal | ACESB | dry | Feb | 2022 | 1/1 (100) |
| Snowy-crowned Robin Chat | ACESB, ACESC | wet | Apr | 2022 | 2/4(50) |
| Speckled Mousebird | AEAGU, Ngel-Nyaki | dry, wet | Apr, Mar | 2021- 2022 | 3/8 (38) |
| Village Weaver | ACESB, ACESC, AEAGU | dry, wet | Feb, Apr | 2021- 2022 | 4/21 (19) |
| Violet-backed Starling | Laminga | wet | May | 2021 | 1/1 (100) |
| Yellow-billed Shrike | ACESB | dry | Dec | 2021 | 1/1 (100) |
| Yellow-mantled Widowbird | ACESB | dry | Feb | 2022 | 1/1 (100) |
| Yellow-throated Greenbul | Laminga | wet | May | 2021 | 1/3 (33) |

## Discussion

Here, we set out to study the circulation and transmission of WNV, USUV, and SINV in African resident and Palearctic migratory birds at two overwintering locations in Nigeria. Our study revealed active circulation of USUV and provided evidence for previous WNV and SINV infections in Palearctic migratory and African resident birds in Nigeria.

USUV RNA was detected in both resident and migratory bird species. This likely reflects overlapping ecological niches and exposure to common vector populations at wintering sites in the same region [51]. The detection of USUV RNA in migratory species in our study further suggests the potential of these birds to contribute to long distance dispersal of the virus through migration. Importantly, one of the USUV-positive birds in our study was an Icterine warbler (a Palearctic migrant) which was sampled in April, prior to spring migration (late February to late June as described by Jonzén *et al.* (2007) [52]. Although, we did not confirm the presence of infectious virus through culture, the detection of viral RNA supports the possibility of active or recent infection. An earlier analysis based on species chorotypes that defined the migratory patterns of birds in Africa and Europe already classified the Icterine warbler as a potential vector of WNV across both continents [53]. The detection of viral RNA in the Garden warbler sampled in October of 2024, corresponded to a period upon arrival at wintering location, but whether the individual returned infected or picked up infection upon arrival or along migratory route could not be ascertained. While no sequences were obtained in our study to confirm this for USUV or WNV, our findings align with previous suggestions of flyway-mediated transmission based on phylogenetic and quantitative models [12,53–55].

USUV RNA could not be detected in the swabs, whereas the feathers of some swab-negative birds were positive. This was likely due to a less optimal cold storage at the sampling locations. Prior to shipping, the samples were kept for weeks under +4°C and then -80°C, which may have degraded the quality of viral RNA if present in the samples. Long persistence of viral RNA in feathers of infected birds has previously been documented for USUV and Influenza viruses [38,56]. Our detections in feathers and not swabs, further emphasized that the collection of feathers may be a promising alternative sample type for virus detection especially under remote field conditions where sustained cold storage cannot be guaranteed.

Antibodies to both WNV and USUV were also detected in resident and migratory birds, with no significant difference in prevalence or antibody titers, suggesting comparable exposure risk across both groups. The concurrent detection of viral RNA and neutralizing antibodies in African thrushes indicates their potential role as reservoirs for USUV, and possibly WNV, in the region. USUV was previously identified in a South African thrush species -a Kurrichane thrush (*Turdus libonyana*) – which was reported to have been sampled in Nigeria [36]. Our findings in African thrushes (*Turdus pelios*) further supports the susceptibility of thrushes to infection in the region. Susceptible host species often share ecological niches [57]. The Song thrush (*Turdus philomelos*), a closely related species in Europe, occupies a similar ecological niche and has repeatedly been reported as a susceptible host of USUV [58,59]. This parallel suggests that members of the genus Turdus may share ecological and biological traits that are relevant for USUV maintenance, although the specific reservoir potential of African thrushes requires further investigation.

Evidence of SINV infection was detected through a screening PCR in a spotted flycatcher, a migratory bird; however, this could not be confirmed by a subsequent confirmatory PCR. Nevertheless, a very recent study at the same sampling location - Amurum, detected SINV in African thrushes [27], confirming enzootic circulation of the virus and the risk of infection to migratory species overwintering at the location. In addition, we found that antibodies to SINV were widely detected across several African resident and European migratory birds in the region, including three out of eight spotted flycatchers. Similarly, SINV antibodies were found in other migratory birds sampled at Amurum. Moreso, antibodies to SINV were detected in migratory and resident birds sampled at Ngel-Nyaki, a location about 300 kilometres away from Amurum, suggesting a broader geographic risk for SINV in Nigeria. Since we neither tested nor compared antibody cross-reactivity to other alphaviruses within the Sindbis virus antigenic group, such as Babanki virus, and to a lesser extent, Semliki Forest virus [60,61], the possibility of mixed infections or misdiagnosis due to cross-reactivity cannot be entirely ruled-out. In the SINV antibody positive migratory birds, the presence of viral RNA could not be confirmed, thus limiting our ability to infer the true source of infections or determine if these individuals were infected at their wintering locations or elsewhere.

## Conclusion

In summary, our findings indicate that migratory birds can become infected with USUV, and potentially with SINV during their overwintering periods in sub-Saharan Africa. Although we did not detect viral RNA in swabs or blood samples, the molecular detection of USUV in feathers suggests possible infections of migratory birds at their wintering locations. Due to poor quality of sera, most of the ELISA positive samples could not be read out in the neutralization assays for functional antibodies, possibly resulting in an underestimation of the number of WNV and USUV infected birds at the study locations. Also, only the ELISA negative samples were tested for SINV, lowering the sample size for our SINV antibody survey. The molecular evidence for USUV, and exposures to WNV and SINV in wild birds in this study, coupled with earlier published molecular evidence for circulation of SINV at one of our sampled locations [27], highlights a potentially wider arbovirus risk in Nigeria that warrants surveillance to understand the epidemiological patterns and public health risks due to possible spillovers. In addition, real-time testing of birds in the region could also maximize the potentials of detecting ongoing infections and aid the recovery of sufficient viral genetic material to sequence and characterize viral strains and provide strong evidence for viral transmission across bird flyways.

## Acknowledgements

We are grateful to Jenny Hesson (Uppsala University, Uppsala, Sweden) for providing the SINV stock and Olli Vapalahti (Helsinki University Hospital, Finland) for the positive controls. We thank Romée van der Beek and Jeroen Vink for their assistance in the laboratory testing of blood and swab samples, and Adams Chaskda, director of APLORI for his support and permissions during sampling. Special thanks to all APLORI CES field assistants who helped during fieldwork.

## Author Contributions

N.C.A: conceptualized the study and wrote original draft. N.C.A, R.S.S and M.P.G.K designed the methodology. J.A.I, N.C.A and Y.B conducted field investigation. N.C.A, A.v.d.L, F.D.C, and C.M.H performed laboratory investigations. N.C.A analyzed and visualized the data. All authors reviewed the manuscript.

